# Optimizing PSC culture for generating neural organoids

**DOI:** 10.1101/2024.11.20.624444

**Authors:** Magdalena A. Sutcliffe, Pia Jensen, Joycelyn Tan, Charles A.J. Morris, Daniel Fazakerley, Martin R. Larsen, Madeline A. Lancaster

## Abstract

Cerebral organoids generated according to unguided protocols produce neural tissue with exceptional cell diversity and fidelity to in vivo. However, with only minimal extrinsic intervention, the importance of high quality starting material becomes paramount. Better understanding of what constitutes a high quality stem cell line and how to maintain those properties throughout prolonged culture is therefore a crucial foundation for successful organoid differentiation. In this study, we investigate the proteome and phospho-proteome of human pluripotent stem cells to uncover the mechanisms that drive neural organoid competence. We identify aberrant cell-extracellular matrix interaction and increased oxidative metabolism as hallmarks of poor neural differentiators. Drawing on the proteomic data and published literature, we optimise culturing conditions by using improved coating matrix, sustained supply of the key growth factor FGF2 and reducing oxidative stress. These adjustments improve brain organoid generation across all tested cell lines, though with varying degrees of efficiency. This work highlights the importance of optimal culture conditions to best support stem cells, ultimately enhancing the quality of brain organoids produced.

## Introduction

In the process of gastrulation, a mammalian embryo generates all major primordia that will later generate all tissue types of the fully formed body. This ability is retained in pluripotent cells cultured in vitro, so called embryonic stem (ES) cells.^1^ ES cells from mouse, human, and other primates have been used to generate a wide array of tissue and cell types, using observations from in vivo to guide the choice of signalling molecules to direct such identities. In vivo and in vitro observations have demonstrated that neural identity determination is the default, and just like in the developing embryo, in the absence of patterning signals, ES cells cultured in vitro naturally adopt a neural fate upon pluripotency exit.^2–4^

The development of technology to produce induced pluripotent stem cells (iPSCs), an ES equivalent reprogrammed from somatic cells, has further added to the strength of in vitro models.^5^ It is now possible to produce PSCs from various healthy and disease-affected individuals through reprogramming. This has resulted in a wealth of rigorously developed and well-characterized iPSC lines, available for researchers to generate specific cell types or tissues of interest from various repositories.^6^

When cultured under optimal 3D conditions without growth factors, ES or iPS cells (collectively termed pluripotent stem cells – PSCs) can form cerebral organoids.^7^ These unguided organoids predominantly exhibit cortical identities, along with adjacent structures such as the hem, choroid plexus, and retina, arranged in a spatially and temporally relevant manner akin to the developing human brain.^8^ Organoids are particularly valuable for modelling development and disease because they provide wider tissue context and faithfully recapitulate natural development and physiology on a micro scale. The application of single cell and omics techniques has allowed organoids to be benchmarked against in vivo tissues and has validated organoids as tools for studying biological processes that were previously inaccessible due to technical or ethical limitations.^9–12^ However, 3D differentiation processes can be sensitive to variability among individual cell lines, especially in methods that rely on natural developmental progression with minimal or no patterning signals. Previous work on brain organoids has observed that the success of the unguided method depends on the physiological state of the pluripotent stem cells, making it highly sensitive to the quality of the starting material.^13–17^

Several research groups have successfully added to our understanding of the variability between PSC lines and provided guidelines on selecting the most suitable cell lines for generating specific tissue types.^18–21^ Jerber and colleagues provided a comprehensive analysis of neural differentiation capacity by characterizing transcriptomic differences between iPS cell lines.^22^ This research evaluated cell lines in the HipSci collection, offering a valuable resource for selecting suitable cells for neural differentiation and identifying markers of poor differentiators that could potentially be applied to independently derived iPSCs. However, the questions of what cellular processes might be responsible for the impaired neural differentiation competence, or whether these differences could be minimized through optimized culture conditions, remain open.

Here, we focus on the cellular proteome and phospho-proteome as indicators of the cellular processes that underlie neural organoid competence. By analysing these proteomic profiles, we aim to optimize the culture conditions of pluripotent stem cells to promote and maintain a state conducive to cerebral organoid formation. This approach provides a deeper understanding of the molecular mechanisms involved and guides the refinement of culturing techniques to enhance organoid development.

## Results

### Proteomic characterisation of organoid competency

To investigate why different cultures show differences in their capacity to differentiate into neural organoids, we selected a panel of six cell lines, comprising three organoid-competent lines (H9, H1, and kolf2) and three lines (sojd3, burb1, and fiaj1) previously demonstrated to be incompetent.^22^ Each group included both male and female lines to account for differences due to sex chromosome makeup. All cell lines were cultured in identical conditions, namely Essential 8 medium on vitronectin (VTN)- coated plates to maintain a fully defined culture environment.

We differentiated each line using the Stemdiff cerebral organoid kit. At the point of organoid generation, we saved 60% of the dissociated cells for proteomic analysis and used the remaining material to produce embryoid bodies (Figure 1A). We included four biological repeats for each cell line. We then monitored the development of the organoids to confirm their competent or non-competent phenotype, allowing us to trace back the organoid differentiation outcomes to the initial cellular material.

**Figure 1.**
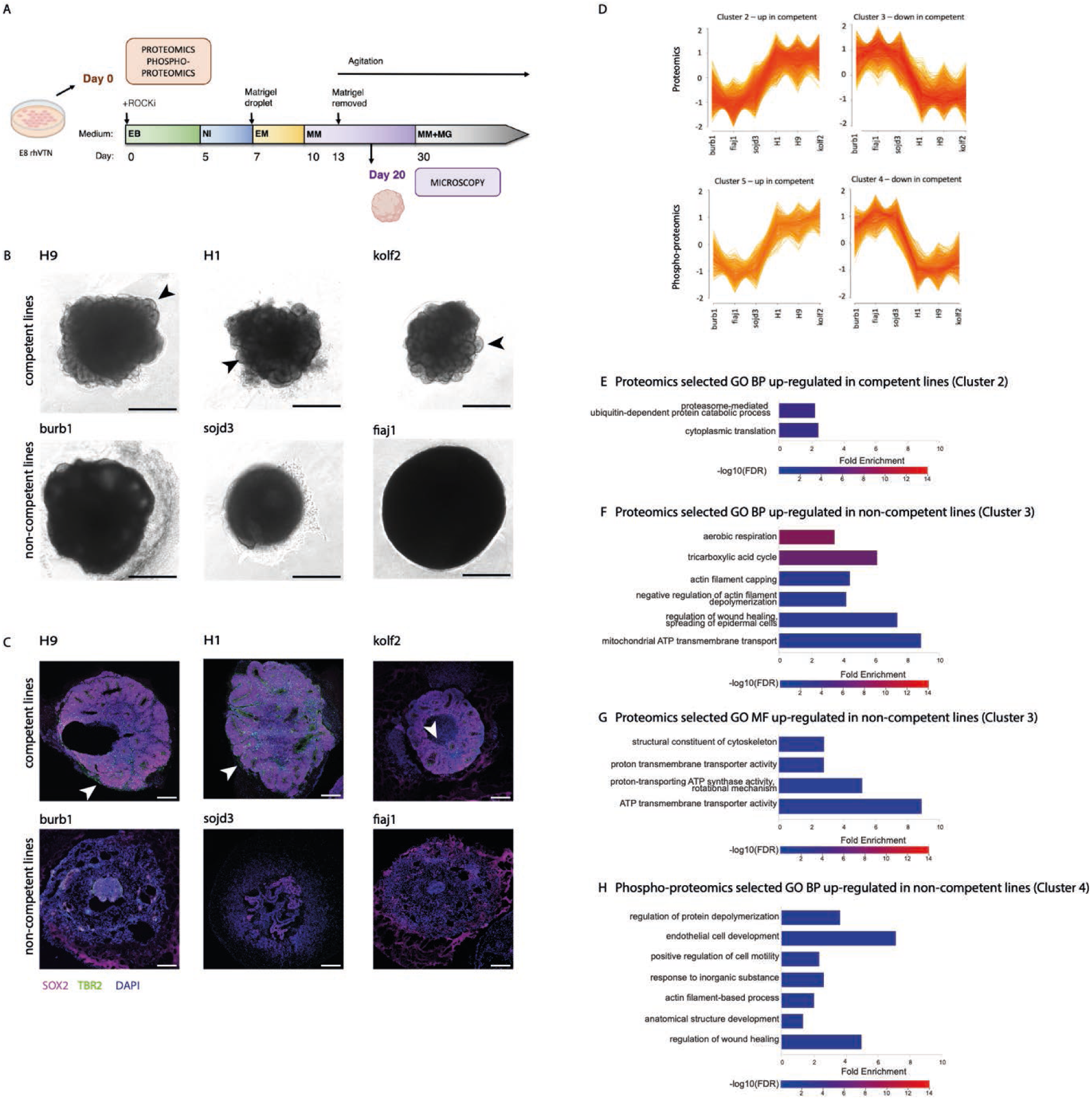
A – schematic of the unguided brain organoid protocol used in this study with indicated sample collections for proteomics and phospho-proteomics, and microscopy, B – day 10 organoids from cell lines used in this study, morphology under phase contrast, black arrowheads indicate neuroepithelial buds, scale bar 500 μm, C – fluorescent images of tissue sections from day 20 organoids made from cell lines used for this study, white arrowheads indicate TBR2+ neuroepithelial buds, D – VSClust analysis - selected clusters that distinguish competent and non-competent cells, top panel – proteomics results, bottom panel – phospho-proteomics results, E- selected GO BP terms overrepresented in cluster 2 – proteins higher in competent cells, F - selected GO BP terms overrepresented in cluster 3 – proteins higher in non-competent cells, G - selected GO BP terms overrepresented in cluster 5 – phospho-proteins higher in competent cells, H - selected GO BP terms overrepresented in cluster 4 – phospho-proteins higher in competent cells, pseudostratified epithelial structures and typical TBR2+ staining (Figure 1C). Based on the day 20 scores, we classified the stem cell lines as either competent or non-competent (Table S1).

Differences in organoid morphology were apparent as early as day 10. Competent cell lines produced organoids with well-defined neural buds, while non-competent lines formed dark structures lacking surface features indicative of neural structures such as inflection points, and instead occasionally developed less cell-dense areas that later became cysts (Figure 1B). Organoids exhibiting correct morphology at day 10 typically progressed to high-quality mature organoids, whereas those with poor initial morphology either failed to improve or deteriorated over time. This was consistent with our previous observations that early organoid morphology reliably predicts mature organoid quality.^23^

We assigned a final organoid quality score at day 20, coinciding with the expression of the dorsal forebrain marker TBR2 (Figure 1C). Cell lines were classified as competent if they produced well-structured organoids with cortical buds comprising pseudostratified SOX2+ progenitors and a visible, scattered layer of dorsal forebrain-specific TBR2+ intermediate progenitors in at least some regions. Conversely, non-competent cell lines produced tissues without visible neural buds, lacking SOX2+

The material saved at the embryoid body generation step was frozen and later processed for proteomics and phospho-proteomics analysis. We conducted a quantitative analysis using tandem mass tag (TMT) LC-MSMS across two separate TMT experiments. This approach identified 7,792 proteins, of which 5,798 identified with two or more unique peptides were used for further analysis (Table S2), and 23,220 phosphopeptides, of which 13,697 were detected in all samples and were used for further analysis (Table S3). Dimensionality reduction was used to examine how the samples grouped together. The 4 repeats for each cell lines clustered very closely together both for proteomics and phosphor-proteomic data. As expected, proteomic profiles of the competent lines H9 and H1 clustered closely, followed by kolf2. The non-competent lines fiaj1 and burb1 also clustered together, whereas sojd3 samples clustered separately (Figure 1S A-B). Interestingly, in the phospho-proteomics analysis, sojd3 clustered with the competent lines H9, H1, and kolf2, whereas fiaj1 and burb1 clustered separately (Figure 1S C-D).

Further cluster analysis using Variance Sensitive Fuzzy Clustering (VSClust) of the proteome data yielded four clusters (Table S4). Cluster 2 was overrepresented in the competent lines, while cluster 3 was overrepresented in the non-competent lines (Figure 1E). We then analysed hits from clusters 2 (upregulated in competent cells) and 3 (upregulated in non-competent cells) for enrichment of GO biological processes (BP) and molecular functions (MF). Cluster 2 showed only two processes enriched more than twofold (Figure 1F) and no statistically significant results for GO molecular function. In contrast, cluster 3 revealed several terms enriched for the TCA cycle, aerobic respiration, and the actin cytoskeleton (Figures 1G and 1H).

An analogous analysis of phospho-proteome resulted in six clusters (Table S5). Hits in cluster 5 were overrepresented in competent cells, while cluster 4 was overrepresented in non-competent cells. Panther GO term analysis did not yield any significant results for biological processes or molecular functions in cluster 5. However, cluster 4 showed one enriched molecular function (cell adhesion molecule binding) and several enriched biological process terms (Figure 1H), primarily pointing to the organization of the actin cytoskeleton.

Taken together, these findings suggest that non-competent cell lines upregulate the actin cytoskeleton and oxidative respiration, providing insights into the molecular underpinnings of cerebral organoid competence.

### Altered actin cytoskeleton and focal adhesions, and the influence of the substrate

We then proceeded to validate the proteomic targets. We stained the actin cytoskeleton in one competent cell line (H9) and one non-competent cell line (fiaj1) to compare their structures. Fiaj1 exhibited thicker actin fibers on the colony edge and in the few peripheral rows of cells, as well as some internal fibres (Figure 2A). In contrast, H9 showed some peripheral actin fibers, but the interior of the colonies displayed a more disorganized, mesh-like actin structure. Actin fibers in pluripotent stem cell colonies are typically associated with focal adhesions (FAs), where cells anchor to the extracellular matrix (ECM). Activated focal adhesion kinase accumulates in focal adhesions.^24^ We stained for active phosphorylated focal adhesion kinase (pFAK-Y397) to determine any differences in focal adhesions between competent and non-competent lines. Indeed, fiaj1 displayed larger and more abundant FAs compared to H9 (Figure 2B).

**Figure 2.**
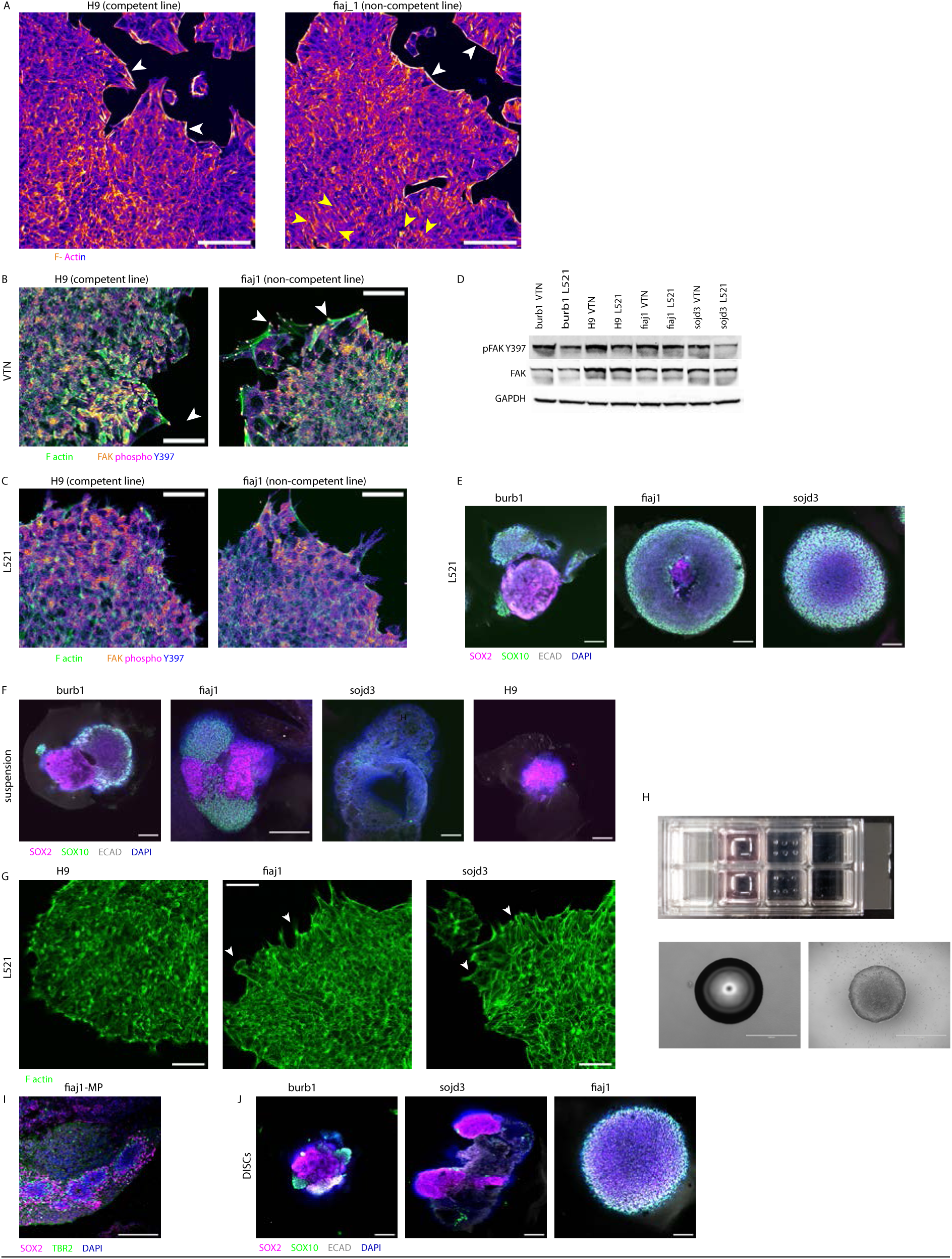
A – phalloidin staining of F-actin in fiaj1 and H9 cells cultured on VTN, white arrowheads indicate peripheral actin fibres, yellow arrowheads indicate actin fibres inside a colony, scale bars 100μm, B – active focal adhesion kinase and actin staining of fiaj1 and H9 cultured on VTN, white arrowheads indicate actin fibres and associated focal adhesions, scale bars 50μm C – active focal adhesion kinase and actin staining of fiaj1 and H9 cultured on L521, scale bars 50μm, D – Western blots of active focal adhesion kinase in sample pairs from 4 lines tested cultured either on VTN or L521, E – day 10 organoids grown from cell lines cultured on L521, scale bars 200μm, F – day 10 organoids grown from cell lines cultured in suspension culture, scale bars 200μm, G – phalloidin staining of F-actin in cell lines cultured on L521, white arrowheads indicate peripheral cell spreading, scale bars 50μm, H – preparation of micropattern colonies – top - droplets of L521 on a 8-well imaging slide, bottom left - a droplet of L521 under phase contrast, bottom right - an attached micropatterned colony, scale bars 1000μm, I – an area of day20 fiaj1 organoids with morphology typical of a competent line, scale bar 200μm, J – day 10 organoids grown from cells cultured on L521 with FGF2 DISCs, scale bars 200μm

Focal adhesions form where integrins interact with the ECM, triggering intracellular complex assembly and signaling. Vitronectin and fibronectin, which engage integrin αvβ5, have been shown to promote the formation of focal adhesions and actin stress fibers in pluripotent stem cells.^24,25^ The peripheral adhesions, so called “cornerstone adhesions” are necessary for maintaining pluripotency but our proteomics and immunofluorescence results suggested that FAs might be overabundant or overly active in non-competent lines. To further investigate this and manipulate actin fibers and focal adhesions, we experimented with different defined substrates. Laminin a5 is abundant in the developing embryonic epiblast, and laminins 521 (L521) and 511 are thought to best support an epiblast-like state in pluripotent stem cells.^26,27^ L521 mainly engages integrin α6β1, which prevents overactivation of FAK that leads to differentiation.^28^

When cells were plated on L521, we observed a reduction in focal adhesion staining. Instead, we saw diffuse FAK staining and the disappearance of thick actin fibers, replaced by a loose actin meshwork with sparse fibers localized to the colony periphery (Figure 2C). To confirm the reduction in FAK activity, we performed a Western blot on cells cultured on VTN versus L521. Most cell lines showed a reduction in FAK phosphorylation when cultured on L521, although it was less pronounced in fiaj1 (Figure 2D).

We then cultured cells on L521-coated dishes to produce cerebral organoids. Organoids grown from burb1 cells cultured in this manner showed slight morphological improvements with large, organised areas showing SOX2+ neural progenitors and with only a small portion of SOX10+ (neural crest marker) disorganised tissue (Figure 2E). However, no improvement was seen in organoids from fiaj1 or sojd3 cells, which consisted mostly of SOX10+ cells and did not form neural buds.

We speculated that ECM interaction might be dysregulated in cell lines unresponsive to the switch to L521, potentially hindering neural differentiation. We adapted the stem cells to culture without any ECM by using suspension conditions as described.^29^ Under these conditions, cells were dissociated and grown with ROCK inhibitor until they formed small clumps that secreted their own ECM. Our proteomics data confirmed endogenous expression of laminin subunits LAMA1, LAMA5, LAMB1, LAMB2, and LAMC5 in standard conditions, in line with previous reports.^30,31^

Suspension conditions did not further improve burb1, which showed similar proportions of SOX2+ and SOX10+ cells in the organoids made from cells cultured on L521 (Figure 2F). Fiaj1 cells grown in suspension produced organoids improved in terms of identity but not morphology; there were more SOX2+ cells but they did not organise in distinct neural buds. Finally, we did not observe improvement in sojd3. Surprisingly, suspension culture in E8 medium led to deterioration of the competent H9 line. H9 organoids grew smaller and although they were positive for SOX2, they did not show typical organisation of neural buds. This suggests that cell-ECM interaction is necessary even in naturally competent cells, and although cells can secrete their own ECM, the concentration might not be sufficient to support stemness.

Since we determined that adherent culture on ECM is necessary but switching to L521 did not improve organoid quality in some non-competent lines, we speculated that aberrant ECM interaction in those cell lines might be the cause. We observed that even on L521, sojd3 and fiaj1 lines tended to spread at the colony periphery, and cells at the edges were less compact than those at the center (Figure 2G). Peripheral cell spreading makes cells more responsive to endogenous growth factors that prevent neural differentiation.^32,33^ We hypothesized that restricting cell spreading could make colonies more uniform in their response to differentiation, as previously demonstrated in 2D models of gastrulation.^34,35^

To test this, we prepared geometrically constrained surfaces for cell attachment on glass slides coated with L521 by applying droplets of L521 mixed 1:1 with PBS containing calcium and magnesium. Gentle pipetting of a 0.5 µL portion of this solution produced a droplet of approximately 1 mm in diameter, forming an adhesive field for cells to attach (Figure 2H). When single fiaj1 cells were seeded on such surfaces, they attached firmly only to the L521 micropatterns, and cells loosely attached to uncoated glass were washed off during medium change (Figure 2H). These micropatterned colonies were then harvested and processed to prepare embryoid bodies for the cerebral organoid protocol. Immunostaining of these organoids revealed regions with correct morphology and marker expression (SOX2 and TBR2), which had not been observed in this cell line before (Figure 2I). However, the overall morphology of the organoids was not improved (Figure S2).

Taken together, these data point to the important role of cell substrate and colony morphology in influencing pluripotent stem cell differentiation capacity. We identified improved results with L521 and micropatterned colonies for certain cell lines, while VTN or a lack of exogenous ECM altogether negatively impacted differentiation capacity.

### Media composition and growth factor signaling

In addition to the substrate, another potential influence on pluripotent stem cell spreading and morphology is media formulation. FGF2 and TGF-β1 in E8 medium are crucial for maintaining stem cell pluripotency,^36^ but FGF2 also plays a significant role in colony morphology.^37^ Although increasing the FGF2 concentration in the culture media does not directly correlate with cell proliferation, it promotes colony compaction and reduces peripheral spreading. E8 medium already contains a high level of FGF2 (100 ng/mL)^38^; however, due to its thermal instability, FGF2 levels decrease by approximately half within the first 4 hours of culture.^39^ To address this problem, we combined L521 coating with a sustained-release FGF2 system using FGF2 DISCs.^40^ This system provides a continuous release of 10 ng/mL FGF2 and requires media changes every 2-3 days with weekly passaging. We switched our non-competent cell lines to L521/FGF2 DISCs in Essential 8 medium for at least two weeks and then used these cells to generate organoids, which were then analysed on day 10.

Burb1 cells cultured under this regimen produced organoids with extensive areas of correct morphology and a small proportion of SOX10+ cells (Figure 2J). Sojd3 also generated organoids with more SOX2+ cells and localized areas of well-structured buds, though these organoids predominantly contained contaminating tissue types that didn’t express SOX10. On the other hand, fiaj1 did not exhibit any improvement under these conditions and displayed the usual rounded morphology with mostly SOX10+ cells. In summary, incorporating sustained-release FGF2 with L521 coating led to improved morphology in burb1 and sojd3 organoids, but did not benefit fiaj1.

### Elevated oxidative metabolism in non-competent lines

Another significant finding from our proteomic screen was the upregulation of oxidative metabolism in non-competent cells (Figures 1G and 1H). Human pluripotent stem cells primarily rely on glycolysis for energy production, converting large quantities of glucose to pyruvate and then to lactate.^41^ In PSCs, the activity of mitochondrial complex I is low, and oxidative phosphorylation (OxPhos) is suppressed, which limits ROS generation and supports genetic stability.^42^ Moreover, glycolysis regenerates NAD+ and provides building blocks for molecules essential for the rapid cell division characteristic of these cells, reviewed in.^43^ Although some pyruvate is directed to the TCA cycle,^44^ the low expression of isocitrate dehydrogenase results in citrate being exported to the cytosol, where it can be converted back to acetyl-CoA by ATP-citrate lyase (ACLY). Acetyl-CoA then serves as a substrate for lipid synthesis and histone acetylation essential for pluripotency maintenance.^45,46^

Although the framework of high glycolysis and low OxPhos is intrinsic to primed pluripotency, metabolism can be significantly influenced by culture conditions. For instance supplying additional glutamine or pyruvate, as well as high insulin can shift the balance away from glycolysis to the TCA cycle and OxPhos.^47,48^ Excess glucose in culture medium leads to increased respiration rates and results in higher peroxide generation.^49^ Lipid deprivation has also been shown to promote TCA and OxPhos.^50^ All these conditions are relevant for culture in E8, which is lipid free but contains glutamine, pyruvate and a high level of insulin and glucose.^38^

Our proteomic data showed higher expression of pyruvate dehydrogenase E1 subunit beta (PDHB) in non-competent lines suggesting that more pyruvate is directed towards the TCA cycle. Therefore, we assessed the conversion of glucose to lactate, a key indicator of glycolytic activity. Typically, glycolysis converts one molecule of glucose into two molecules of lactate. All cell lines exhibited robust conversion of glucose to lactate, with a slightly higher glucose-to-lactate ratio in the competent H9 line (Figure 3A) suggesting a greater reliance on glycolysis than the other cell lines.

**Figure 3.**
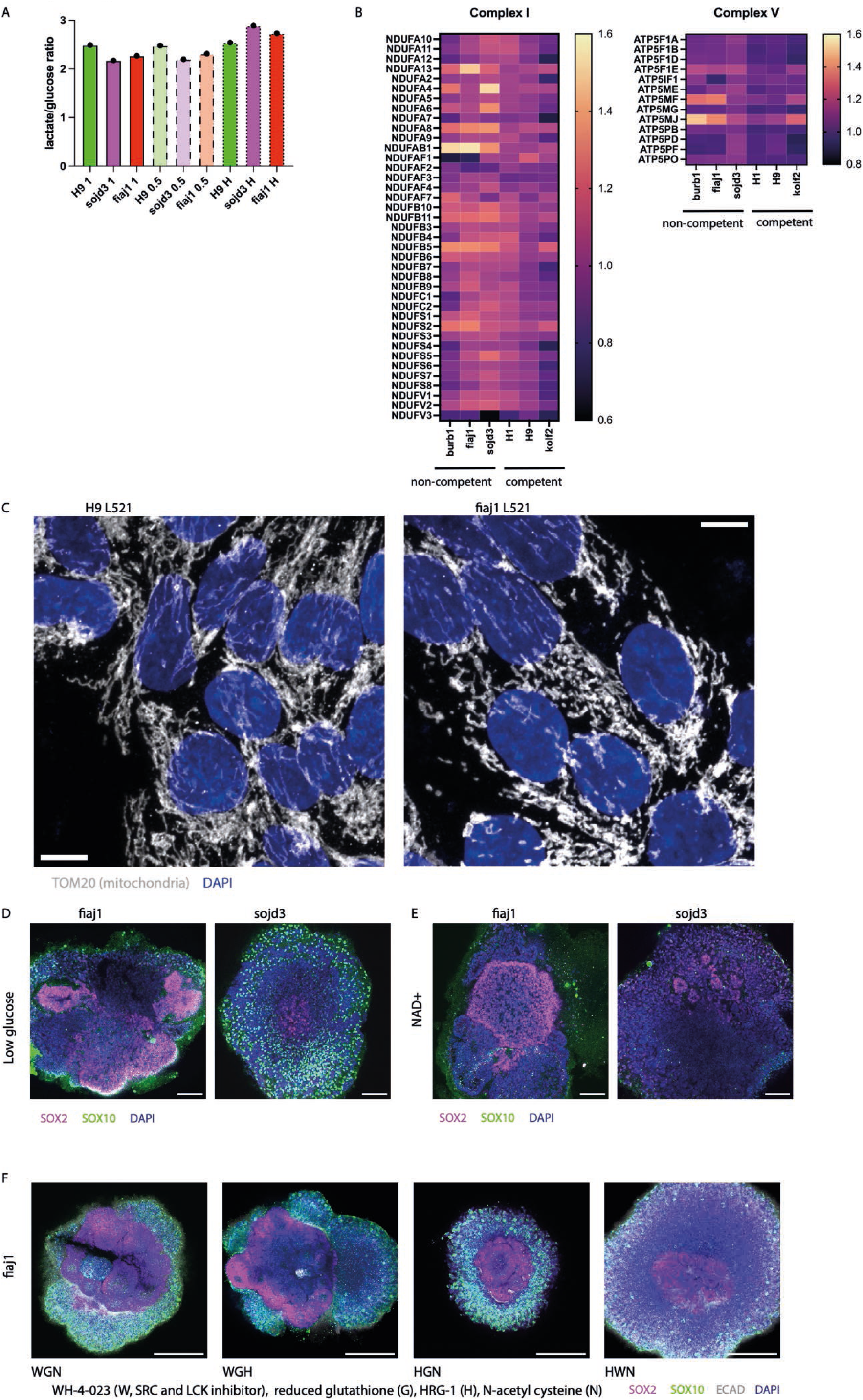
A – ratio of conversion of glucose to lactate in one competent (H9) and two non-competent lines (fiaj1, sojd3) in 1ml of medium (1), 0.5ml of medium (0.5) or in 0.5ml under hypoxia 5% oxygen (H), B – heatmaps of protein abundances of all detected component of mitochondrial electron transport chain complexes 1 and 5, C – mitochondrial morphology in a competent (H9) and non-competent (fiaj1) cell line cultured on L521, scale bars 10μm, D – day 10 organoids grown from cells cultured in low glucose conditions, scale bars 100μm, E - day 10 organoids grown from cells cultured in with 50μM NAD+, scale bars 100μm, F- day 10 organoids grown from cells cultured with combinations of (W) WH-4-023, SRC inhibitor, (H) heregulin beta, (G) reduced glutathione and (N) N-acetyl cysteine, scale bars 200μm

In addition, cellular metabolism is affected by oxygen availability particularly in cells dependent on OxPhos. Cells cultured under atmospheric oxygen concentrations under a thick layer of medium can lead to local hypoxia due to limited diffusion.^51^ We therefore also tested cultured cells in either half-volume media, or half-volume media under hypoxic conditions (5% O2) compared to normal volume of media (600ul in 1 well of 24-well plate), and measured glucose and lactate levels after 24 hours. Conversion rates were similar in both full and half-volume media, suggesting that oxygen diffusion at atmospheric concentration is not a limiting factor driving metabolic shifts between OxPhos and glycolysis. In hypoxic conditions, the competent cell line maintained its glucose-to-lactate conversion ratio, while the non-competent lines showed an increase, indicating that these suboptimal lines might be more influenced by environmental oxygen levels (Figure 3A). This is consistent with a slightly increased reliance on OxPhos seen in the proteome of non-competent cells (Figure 1F), and therefore a potential greater sensitivity to oxygen availability.

Interestingly, in all PSC lines tested, the ratio of lactate produced to glucose consumed exceeded the expected value of 2, suggesting that pluripotent stem cells utilize additional substrates from the culture media to produce lactate. Notably, the amount of glucose consumed from the medium was only a small fraction of the total glucose concentration available, implying that with daily medium changes, cells are exposed to unnecessarily high glucose levels (Figure S3).

We observed upregulation of multiple components of complexes 1 and 5 of mitochondrial electron transport chain in non-competent lines (Figure 3B) further suggesting more active OxPhos. This is atypical of pluripotent stem cells, which normally downregulate activity of complex 1 and suggests differences in mitochondrial physiology between competent and non-competent lines.^42^ To confirm this, we assessed mitochondria morphology in the competent H9 line and the non-competent Fiaj1 line, both cultured on L521. In fiaj1 cells, mitochondria appeared smaller and more dispersed, whereas in H9 cells, they were elongated and interconnected (Figure 3C).

Whilst OxPhos is superior to glycolysis in terms of ATP generation, glycolysis plays a critical role in regenerating NAD+ for various metabolic processes and generating building blocks in rapidly proliferating pluripotent stem cells.^52^ At the same time directing pyruvate away from lactate generation and into TCA suppresses cell proliferation by reducing the NAD+/NADH ratio. NAD+ supplementation in hESC decreases cellular dependency on glycolysis to maintain favourable NAD+/NADH ratio and improves pluripotency marker expression.^53^ Since the non-competent cell lines display reduced glycolysis and increased TCA activity, we hypothesized that they might be depleted in NAD+. We therefore cultured cells in a physiological concentration of glucose or supplemented NAD+ to correct the NAD+/NADH ratio. To do so, we modified the culture medium with glucose concentration adjusted to 5mM from the original 17mM in E8 medium, or treated cells with 50μM NAD+ for at least 3 passages and generated organoids. Day 10 organoids derived from fiaj1 non-competent cells cultured in medium with 5 mM glucose on L521 demonstrated improved marker expression, larger SOX2+ areas, and better morphology (Figure 3D). In contrast, sojd3 cells did not show similar improvements under these conditions. Supplementation with NAD+ also improved SOX2 expression in fiaj1 organoids but did not have a significant effect on sojd3 cells (Figure 3E).

Finally, we explored the role of antioxidants in mitigating the oxidative stress caused by upregulated OxPhos. In a study of cardiomyocyte differentiation, supplementation of antioxidants proved as effective as glucose level reduction in reducing ROS generation and downstream signalling.^49^ To test if addition of antioxidant improved organoid competency, we added 2.5mM of N-acetyl cysteine (NAC) and/or 1mM reduced glutathione (GR) to regular glucose E8 medium. Because of our findings above with stabilized FGF2 and focal adhesions, we also combined these antioxidants during cell culture with heregulin beta 1 (HRG1), which enhances FGF2 action through ERBB2 phosphorylation,^54–56^ and with WH-4-023, a SRC inhibitor (SRCi), during embryoid body (EB) formation. Antioxidants, either with HRG1 or SRCi, were maintained during the first 3 days after EB generation. The combination of the SRC inhibitor and GR resulted in the most significant improvements in marker expression and morphology (Figure 3F), while HRG1 did not enhance these features to the same extent. Additionally, GR was found to be more effective at improving organoid differentiation than NAC.

## Discussion

Optimal cell culture conditions of pluripotent stem cells are crucial for maintenance of the two key features of PSCs: self-renewal and trilineage differentiation potential.^57^ Although human ESCs have been cultured for over 25 years, now, and iPSCs for almost 20 years, we are still learning about aspects of PSC biology, and more work is needed on finetuning conditions to achieve the best possible fidelity to in vivo. Whereas the aspect of self-renewal of PSCs seems to be well understood and commercial media sustain cell division and maintenance of key pluripotency markers, the differentiation potential appears more problematic.^58^ This particularly affects the organoid field, where any small inconsistencies in the starting material are amplified throughout protocol timelines and result in low yields or complete failure of differentiations. Unguided brain organoids seem to be particularly sensitive to the state of PSCs, whereas their correct complex morphology is crucial to faithfully reflect natural brain development and function.^23^

In this study we have shown that organoid competence can be improved by optimised PSC culture conditions. We addressed two areas of cell physiology informed by findings from our proteomic screen, namely dysregulated cell adhesion and aberrant metabolic shift towards OxPhos in non-competent cell lines. We found that an L521 substrate, steady supply of FGF2 and reducing the impact of increased OxPhos were able to increase the proportion of SOX2+ neural tissues and decrease the proportion of other tissues in unguided cerebral organoids.

Recently, two other studies proposed optimised culture conditions that eliminate the need to use feeder cells to achieve improved organoid morphology, also to cell state before pluripotency exit as crucial for cerebral organoid success. ^14,15,59^ They collectively focus on balancing the delicate interplay of PI3K and ERK signalling activated by FGF2 and SMAD activated by TGFbeta ligands, whose cross-talk and relative levels maintain pluripotency or lead to differentiation.^54^ Here, we did not test TGFbeta signalling in PSCs because proteomic results did not highlight it as a potential differentiator between competent and non-competent cell lines, but we found that TGFbeta inhibition at the embryoid body stage led to generation of better tissues (Table 1). We addressed multiple aspects of FGF signalling: pre-treatment with FGFR1 inhibitor or inhibition of the two major effector of FGF-signalling. PI3K inhibition slightly improved organoids, whereas FGFR1 or MEK1/2 inhibition did not (Table 1). We found that a sustained FGF supply is optimal and could prevent downstream pathway activity fluctuations, as well as improving colony morphology, as suggested earlier.^37^ We showed that this effect on cell state and morphology also led to improved cerebral organoids for some cell lines. Although not tested in our work, it might also be possible to achieve a similar effect with engineered thermostable forms of FGF2, which are now commercially available ^60,61^.

**Table 1.**

| Treatment | Rationale and references | Effect | Data shown in this paper |
| --- | --- | --- | --- |
| culture on L521 | L521 engages integrin $\alpha 6$ and prevents overactivation of FAK that leads to loss of pluripotency,<sup>28</sup> cells cultured on laminin show fewer large focal adhesions and fewer actin stress fibres than when cultured on vitronectin, proteomics results showed more dynamic actin cytoskeleton in bad differentiators | improved organoid generation in burb1 | Yes, Figure 2E |
| inhibition of SRC at basal culture (CGP-77675) | Direct inhibition of FAK was avoided due to literature evidence that some activity of FAK is needed for survival and pluripotency maintenance<sup>75</sup> and that FAK plays a role in pro-survival signalling through IGF1R<sup>76</sup> | Worse organoid differentiation outcome, differentiation of PS cells with prolonged exposure | No |
| inhibition of SRC at organoid generation (WH-4-023) | chemical inhibition of SRC family kinases or knockdown of SRC improve differentiation towards all germ layers<sup>77</sup>, inhibition of SRC caused epithelial differentiation<sup>78</sup>, bad differentiators show higher FAK signalling that is improved with L521 | no improvement alone, improves outcome when used alongside with HRG and antioxidants in fiaj1 line | Yes, Figure 3F |
| FGF2 DISCs - sustained FGF2 release, 3 feeds a week | FGF2 is necessary to maintain pluripotency and its withdrawal leads to differentiation<sup>79</sup>, sustained release FGF2 from DISCs improves expression of stem cell markers, increases stem cell numbers and decreases spontaneous differentiation<sup>39</sup> | in cells cultured on L521 improved organoid differentiation in burb1, improved marker expression in sojd3 but with bad morphology, no difference in fiaj1 | Yes, Figure 2J |
| culture with HRG | bFGF, a growth factor essential for PSC maintenance, leads to phosphorylation of ERBB3, a HRG co-receptor<sup>55</sup>, self-renewal of PSCs requires ERBB2/ERBB3 activation that can be achieved with HRG<sup>56</sup>, | some improvement in fiaj1, better if combined with antioxidants and/or SRC inhibitor WH-4-023 | Yes, Figure 3F |
| geometric constriction - culture on L521 micropatterns | culture of PSCs on micropatterns standardises their response to gastrulation inducing cues in vitro<sup>34</sup>, constrained cells in the centre of micropatterns are insensitive to WNT signals that induce gastrulation like process at the colony periphery<sup>80</sup> | Some improvement in fiaj1 (areas of correct differentiation present but a small percentage of overall organoid volume) | Yes, Figures 2J |
| organoid culture with dual SMAD inhibition | Noggin/SB431542 facilitate neural induction in human PSCs<sup>81</sup>, short pulse of Noggin/SB431542 used for 1-3 days in the first 3 days of organoid protocol | Only using Noggin/SB431542 for longer shows any improvement, leading to change of the character of the differentiation protocol from unguided to guided | No |
| generation of organoids with MEK inhibition | addition of PD0325901 (inhibitor of MEK and downstream ERK) improves neural ectoderm differentiation from mouse epiblast stem cells,<sup>82</sup> FGF signalling prevents neural induction through ERK activation in human PSCs<sup>83</sup> | Some improvement in bad batch of kolf2, no improvement in fiaj1 | No |
| resetting cells to naïve by Rset (Stemcell Technologies) and then re-priming in | hypothesis - resetting cells to naïve conditions could erase aberrant epigenetic marks that prevent neural differentiation, done with Rset protocol (Stemcell Technologies) | cells die at EB stage if EBs were generated straight from cells cultured in Rset, when cells were returned to primed conditions organoids form but | No |
| Stemflex (Thermo) |  | with no improvement to morphology in non-competent batch of kolf2 cells |  |
| reducing glucose in culture medium | high levels of glucose in standard culture media for PSCs contribute to oxidative stress and bias cells towards mesoderm and cardiomyocyte differentiation<sup>49</sup> | Lower glucose and no pyruvate improve morphology in fiaj1 cells | Yes, Figure 3D |
| eliminating pyruvate from medium | exogenous pyruvate enhances mesodermal differentiation of PSCs<sup>48</sup> | Lower glucose and no pyruvate improve morphology in fiaj1 cells | Yes, Figure 3D |
| long term culture in E8 medium | culture in E8 leads to improved directed neural differentiation,<sup>50</sup> E8 medium was used as the default condition in this study | no improvement and in some cell lines (H9) deterioration | Yes |
| addition of alpha-ketoglutarate to generate EBs | addition of alpha-ketoglutarate improves differentiation of primed pluripotent stem cells though modification of histone and DNA methylation<sup>84</sup>, culture in E8 leads to improved directed differentiation into neural tissue through increasing alpha-ketoglutarate/succinate ratio<sup>50</sup> | Embryoid bodies don't form due to cell death | No |
| pre-treatment with FGFR1 inhibitor | FGFR1 inhibition for two days before organoid generation improves identity and morphology to the same extent as culture on feders<sup>15</sup> | no improvement, in sojd3 deterioration | No |
| antioxidants (reduced glutathione and NAC) added to PSC culture and for the first 3 days of EB culture | high level of oxidative metabolism and ROS bias towards mesoderm differentiation tha can be reversed by using antioxidants,<sup>48,49</sup> proteomics results suggest increased mitochondrial oxidative metabolism in bad differentiators | some improvement in fiaj1, better when combined with HRG and/or SRC inhibitor WH-4-023 | Yes, Figure 3F |
| inhibition of PI3K during EB generation | PI3K drives PSC survival but also increases oxidative metabolism,<sup>47</sup> there is some phenotypic overlap at transcriptomic and proteomic level between bad cell lines and cells with mutant PIK3CA H1047R allele<sup>85</sup> suggesting potential overactivation of PI3K pathway | addition of PI3K inhibitor at the point of EB generation together with WNT inhibition improve organoid differentiation in fiaj1 line | No |

In this study, we also found that the extracellular matrix substrate that cells are cultured on plays a large role in organoid competency. L521 has been developed to provide fully defined culture conditions while reducing cell death with passaging and single cell cloning.^62^ It has also been demonstrated to improve pluripotency marker expression.^26^ We show that culture on L521 corrected abnormal cytoskeletal morphology and overactive focal adhesions, and improved organoid competence in some cell lines alone and in combination with other interventions in all of them. For these experiments, we did not use Matrigel, a coating matrix frequently used for PSC culture, due to its undefined formulation, more promiscuous integrin binding pattern and potential batch to batch variability^63,64^; however, its laminin-rich nature may explain why it has previously been described to be a successful substrate for PSCs when generating cerebral organoids.^65^ Interestingly, direct inhibition of focal adhesion signalling was detrimental to cell survival (Table 1). Several proteins associated with focal adhesions and actin cytoskeleton were previously shown to be upregulated in iPSC vs ESC, pointing to the sensitivity of these process to in vitro culture conditions.^66^ However, at a global level, differences between ES and iPSC were very minor^67^.

Further, we found that metabolism of pluripotent stem cells also affects organoid competency. Cell lines that showed increased TCA and OxPhos tended to perform worse than cells with more glycolytic metabolism. Similar findings were reported by Yamamoto and colleagues, where a glycolytic shift improved the differentiation potential of H9 ES cells.^68^ Essential 8 medium, which we used due its simple and fully defined composition, tends to promote more TCA activity and oxidative metabolism in PSCs.^50^ Interestingly, Cornacchia and colleagues observed improved directed neural differentiation with E8 medium.^50^ It is possible that this reflects differences between unguided and directed differentiation, similar to the contrasting findings of TGFbeta signalling supporting organoid development in the unguided method vs impeding organoid differentiation when using dual SMAD inhibition methods.^14,40^ It is also possible that non-competent cells may aberrantly upregulate oxidative metabolism in E8 or struggle to manage excess ROS, which could impact their differentiation potential.^49,69^ We also noticed that the glucose concentrations in commercially available stem cell culture media are unnecessarily high and that with daily media changes, PSCs only use up a small fraction of the available glucose. Interestingly, findings of another large proteomic study corroborate our conclusion that culture media and the presence of feeder cells influence the PSC proteome.^70^ Culture conditions were the second strongest influence on protein expression in iPSCs, after genetic background, and affected proteins involved in cholesterol biosynthesis, transcription, translation and vesicular transport.

A recent comparative proteomic study of iPSCs and ESCs pointed towards metabolic differences between the two PSC types.^71^ Comparison with our findings revealed that some similarities exist between the features that distinguish ESC from iPSCs and competent from non-competent cells, but also some important differences. For example, the iPSCs interrogated by Brenes and colleagues upregulated components of all complexes of the electron transport chain. However, the non-competent cell lines used here specifically showed higher abundance of complex 1 and 5 proteins. hiPSCs also showed higher expression of multiple nutrient transporters and enzymes in key metabolic pathways, as well as generally higher protein content per cell. Several of the highlighted metabolic hits also showed downregulation between competent and non-competent lines (CPT1A, SLC25A20, ACO2 IDH3A-G, SDHA-B, OGDH, GLUD, GLS, OAT, GOT2) but some displayed inverse or another pattern (MECR, DLD, GPT2). This suggests that the metabolic changes might not be iPSC-specific but that iPSC lines might have less robust metabolic control and in suboptimal conditions some slip into an elevated metabolic state not compatible with competency.

Finally, we observed that the primary contaminating cell types in our non-competent differentiations were SOX10-positive neural crest-like cells. Neural crest (NC) development occurs at the neural plate border, which is exposed to WNT and BMP signals causing NC cells to delaminate from the neuroepithelium and acquire a migratory phenotype (reviewed in^72^). Sensitivity to WNT and WNT signalling regulation have been reported to influence fate decisions after pluripotency exit,^73,74^ however, our screen did not reveal elevated canonical WNT or BMP signalling. This leads us to speculate that abnormal elevation of BMP and WNT signalling could occur during differentiation rather than during PSC culture.

In conclusion, we demonstrate that optimisation of culture conditions by seeding on L521 substrate, steady supply of FGF2 and lowering OxPhos or addition of antioxidants can improve cerebral organoid differentiation. Although combinations of these changes produced some degree of improvement in each of the cell line tested, we are aware it is not a universal solution and may need to be tailored and tested for individual lines.

## Materials and methods

### Cell lines

Two human ESC lines, and four human iPSC lines were used in this study. The ESC lines were H9 (WA09, female) and H1 (WA01, male) and were purchased from WiCell. The iPSC lines: burb1 (HPSI0714i-burb_1, hPSCreg WTSIi257-A, male), fiaj1 (HPSI0514i-fiaj_1, hPSCreg WTSIi301-A male), kolf2 (HPSI0114i-kolf_2, hPSCreg WTSIi018-B, male) and sojd3 (HPSI0314i-sojd_3, hPSCreg WTSIi073-A, female) were obtained from hispci.org. The use of human ESCs used for this project was approved by the U.K. Stem Cell Bank Steering Committee and iPSCs and ESCs were approved by an ERC ethics committee and are registered on the Human Pluripotent StemCell Registry (hpscreg.eu).

### Cell culture

All cell lines were cultured in Essential 8 (E8) medium (Thermo Fisher Scientific) on TC-treated 6 well plates (Corning) coated with rh-VTN (Thermo Fischer Scientific) at xug/ well. Cultures were split as clumps twice a week using 0.5mM EDTA. Where specified, culture plates were coated with rhL521 at 0.5mg per well in DPBS with calcium and magnesium. For some experiments cells were cultured in medium low glucose low insulin medium consisting of DMEM no glucose no pyruvate (Thermo), Sodium bicarbonate 7.5% Ascorbate Mg phosphate (Merck), B27-A (Thermo Fisher), N2 (Thermo Fisher), MEMNEAA (Merck), Glucose 5mM and growth factors FGF2 (100ng/ml), and TGFb (2ug/ml). This modified medium did not contain pyruvate, contained antioxidants from B27-A (DL Alpha Tocopherol Acetate, DL Alpha-Tocopherol, reduced glutathione, catalase and superoxide dismutase) and contained lower concentrations of insulin (5.3mg/l vs 19.4mg/ml) and glucose (5mM vs 17.5mM) than E8 medium.

### Cerebral organoid generation

Cerebral organoids with telencephalic identity cells were generated as described previously using STEMdiff Cerebral Organoid Kit (StemCell Technologies, 08570), Figure 1A Lancaster et al. 2017). Briefly, cultures at <80% confluence were washed with PBS and dissociated using Accutase. 9,000 cells were seeded in each well of a round bottom ultra-low adhesion 96 well plate (Corning) in EB medium with 10uM Y27632 and left for 3 days to form embryoid bodies. Medium was replaced on day 3 with fresh EB medium without 27632 and then changed to NI on day 5. EBs were embedded in Matrigel (Corning) droplets on day 7 and transferred to Expansion medium. On day 10 droplets were transferred to Maturation Medium. On day 13 Matrigel was removed using mechanical Dissociation and/or 30 minute incubation with Cell Recovery Solution (Corning), then tissues were transferred to fresh Maturation Medium and cultured with agitation.

### Histological and immunohistochemical analysis

Samples were fixed in 4% PFA either overnight at 4 °C or at room temperature for 1h, then washed twice in PBS for 10 min. Samples for cryosectioning were incubated overnight in 30% sucrose in 0.2M PB (21.8 g/l Na2HPO4, 6.4 g/l NaH2PO4 in dH2O), embedded in gelatin (7.5% gelatin, 10% sucrose in 0.2 M PB) and plunge frozen in 2-methylbutane (Sigma-Aldrich, M32631) at below −30°C. Frozen blocks were sectioned at the thickness of 20 μm and stained as previously described (Lancaster et al., 2013). Wholemount samples were stained for 24-48h with primary antibodies followed by 24-48 h with secondary antibodies in a buffer of 4% normal donkey serum and 0.25% Trton-X-100 in PBS.

### Antibodies and stains

Primary antibodies used in this study were as follows: SOX2 (Abcam, ab97959, 1:200 for IF), TBR2 (R&D Systems, AF6166 1:200 for IF), HuC/D (Invitrogen, A2127, 1:200 for IF), FAK (Abcam, ab40794, 1:1000 for WB), pFAKY397 (Thermo Fisher Scientific, 44-624G, 1:200 for IF, 1:1000 for WB), GAPDH (Abcam, ab8245, 1:5000 for WB), SOX10 (R&D Systems, AF2864, 1:100-1:200 for IF), ECAD (BD Transduction, 610181, 1:400 for IF), TOM20 (Santa Cruz, sc-17764 1:500 for IF). Secondary antibodies used were: Donkey a-Ms 488 (Life Technologies, A21202), Donkey a-Rb 488 (Life Technologies, A21206), Donkey a-Sh 488 (Life Technologies, A11015), Donkey a-Ms 568 (Life Technologies, A10037), Donkey a-Rb 568 (Life Technologies, A10042), Donkey a-Ms 647 (Life Technologies, A31571), Donkey a-Rb 647 (Life Technologies, A31573), Donkey a-Sh 647 (Life Technologies, A21448). All secondary antibodies were used at 1:500 dilution. Nuclei were counterstained with 0.1 µg/ml DAPI (Merck, 268298). F-actin was detected with ActinGreen 488 ReadyProbes Reagent (Alexa fluor 488 conjugated phalloidin, Invitrogen).

### Imaging and image analysis

Images were acquired on Zeiss LSM 710 or Zeiss LSM 780 systems with each channel as separate track. Images were acquired at x100, x200 or x630 magnification. Raw images were processed using FIJI and brightness and/or contrast were adjusted where needed for clarity.

### Protein extraction, digestion and Tandem Mass Tag labeling

At the point of organoid generation, 60% of Accutase dissociated cells were pelleted at 200rcf, supernatant removed and remaining pellets frozen at −70°C. Material was collected from four independent batches for each cell line. Cell pellets were dissolved in 150 µl lysis buffer consisting of 1% (w/v) sodium deoxycholate (SDC, Sigma), 10 mM Dithiothreitol (DTT, Sigma) and 50 mM triethylammonium bicarbonate (TEAB, Sigma), pH 8 and sonicated on ice for 4x 10s using a probe sonicator at 40% amplitude. Protein concentrations were measured using a NanoPhotometer® N60 (Implen) against a standard curve of HeLa cell protein extract. 50 µg of each sample was alkylated with 20 mM Iodoacetamide (IAA, Sigma) for 30 minutes in the dark followed by digestion with 5% (w/w) in-house methylated trypsin (Sigma) (Heissel et al. 2018; https://doi.org/10.1016/j.pep.2018.03.002) for 4h at 37°C.

Thereafter the peptide samples were labeled with Tandem Mass Tag (TMT) 16plex Isobaric label Reagents (Thermo Scientific) according to the manufacturer’s instructions. Two sets of a TMTpro 16-plex were used and in each TMT set, one channel was used for labelling of a pooled sample to enable comparison of samples between the TMT sets. The labelling reaction was checked by LC-MS/MS analysis to ensure proper labelling of all TMT channels and excess reagent was quenched using 5% hydroxylamine (v/v) (Thermo Scientific) for 15 min at RT. After incubation, the labeled peptides were mixed 1:1 and the pooled TMT samples were acidified with 2% Formic acid (FA) and vortexed to pellet SDC. The samples were centrifuged at 20,000 xg for 15 min at RT and the supernatant was transferred to a new tube and dried by vacuum centrifugation.

### Enrichment of phosphorylated peptides

The TMT labelled peptide mixture was dissolved in 80% acetonitrile (ACN), 5% trifluoroacetic acid (TFA) with 1 M glycolic acid (Sigma) and incubated with 0.6 mg TiO_2_ beads (Titansphere 10 µm, GL Sciences) per 100 µg peptide for 15 min at RT with vigorous shaking to enrich the phosphorylated peptides ^86^ The beads were centrifuged briefly, and the supernatant transferred to a new tube with 0.3 mg TiO_2_ beads per 100 µg peptide. After 10 min incubation at RT with vigorous shaking and a brief centrifugation the supernatant was collected. The beads were subsequently washed with 80% ACN/1% TFA and 10% ACN/0.1% TFA. The supernatant with the unbound TiO_2_ fraction and the washing fractions, both containing the non-modified peptides, were combined. The phosphorylated peptides were eluted from the beads by incubation with 1.5% ammonium hydroxide solution (Sigma), pH 11.3, at RT with vigorous shaking. The beads were spun down and the supernatant passed through C8 material from a 3M Empore^TM^ disk (Sigma). Any remaining peptides were eluted from the disk with 30% ACN and all peptide samples were dried. Since sialylated glycopeptides also bind to the TiO_2_ beads (Larsen et al. 2007; https://doi.org/10.1074/mcp.m700086-mcp200), the sample was de-glycosylated with N-glycosidase F (Biolabs) and Sialidase A (Prozyme) in 50 mM TEAB, pH7.5 at 37°C ON.

### High-pH fractionation

To reduce the complexity of the samples, non-modified and phosphopeptides were fractionated by High-pH chromatography prior to nanoLC-MS/MS analysis.

The peptide samples were dissolved in 30 μL solvent A (20 mM ammonium formate, pH 9.5) and loaded onto an Acquity UPLC^TM^M-Class CSH^TM^C18 column (Waters) using a Dionex Ultimate 3000 HPLC system (Thermo scientific). Approximately 100 μg of the non-modified peptide samples was fractionated, whereas the whole of the phosphopeptide samples were fractionated. Separation of the peptides were performed using a 70-minute gradient from 2 to 95% solvent B (80% ACN, 20% solvent A) in solvent A at a flow rate of 0.1 μL/min. The fractions are collected every 60 seconds into a final of 12 concatenated fractions in a 96-well plate (Axygen) and subsequently dried by vacuum centrifugation and stored at −20°C.

### Reversed-phase nanoLC-ESI-MS/MS

Each high pH fraction was resuspended by adding 3 μL of solvent A (0.1% FA) and loaded in a volume of 2.5 μL onto an analytical column on an EASY-nLC 1000 system (Thermo Scientific).

The analytical column was a 21 cm long fused silica capillary (75 µm inner diameter) and packed with ReproSil-Pur C18 AQ 1.9 µm reversed-phase material (both resins Dr. Maisch Ammerbuch-Entringen). The peptides are eluted with an increasing concentration of organic solvent (solvent B: 95% ACN, 0.1% FA) over a gradient of 120 min in the following manner: from 2% to 25% solvent B in 100 min, 25- 40% in 20 min and 40-95% in 1 min. The flow was 300 nL/min. The nLC was online connected to an Orbitrap Eclipse^TM^ Tribrid^TM^ mass spectrometer (Thermo Scientific) operated at positive ion mode with data-dependent acquisition. The Orbitrap acquire the full MS scan with an automatic gain control (AGC) target value of 300% (3×10^6^) ions and a maximum injection time of 50 ms. Each MS scan is acquired at high-resolution (120,000 full width half maximum (FWHM)) at m/z 200 in the Orbitrap with a mass range of 350-1600 Da. For the non-modified peptides, the peptide fragmentation was performed using the SPS-MS3 method with real time database searching^87^ Briefly, each peptide was selected (0.7Da window) and fragmented in the linear ion-trap using CID with a normalized collision energy of 35% and an activation time of 10 ms. Each MSMS spectrum was subjected to a brief 20-30 ms database search against a Human uniprot reviewed database using a database search program build into the MS computer. If the resulting MSMS database search received a confident match in the database, the same ion was reselected and fragmented in the linear ion-trap and subsequently the 10 most intense fragment ions originating from the identified peptide was reselected and fragmented using HCD fragmentation (NCE 55) and scanned out in the orbitrap with 30.000 in resolution optimized for resolving of the TMT reporter ions. The eclipse workflow was set to automatic calculation of number of peaks that could be selected within a 3 second duty cycle. For the phosphopeptides, the fragmentation was performed using HCD NCE 36, with the following setting for ion detection; resolution 50K FWHM, maximum injection time 200 ms and AGC target 200%. The MSMS was performed with a cycle time of 3 sec. All raw data are viewed in Thermo Xcalibur v3.0.

### Mass spectrometry data analysis

The raw data were processed using Proteome Discoverer (v2.5, ThermoFisher, PD2.5) and all data were searched against a Human uniport reviewed database. The non-modified peptides were searched using the SEQUEST HT search algorithm only whereas the phosphopeptide data were searched first using an in-house Mascot search algorithm followed by the SEQUEST HT search algorithm. The data were searched with 10 ppm accuracy in MS and 0.8 Da in MSMS mode (linear ion trap MSMS) for the non-modified peptides and 0,05 Da for the phosphopeptides. The quantitation was performed from the MS3 HCD spectra for the non-modified peptides and MS2 HCD spectra for the phosphopeptides. Database searches were performed with the following parameters: TMTpro 16-plex (Lys and N-terminal) as fixed modifications and a maximum of 2 missed cleavages for trypsin. In addition for the phosphopeptides, the search was performed with phosphorylation of serine/threonine/tyrosine (S/T/Y) and deamidation of asparagine (N) as variable modifications. All identified peptides were filtered against a Decoy database using Percolator with a false discovery rate (FDR) of 0.01 (FDR < 0.01). Only peptides with rank 1 were considered for further analysis. Only proteins with more than 1 unique peptide were considered for further analysis in the non-modified group.

Quantitative analysis was based on 4 biological replicates. Quantification across the 2 sets of TMTpro 16-plex was normalized based on a common reference channel containing a mix of all samples. The relative abundances of the non-modified and phosphopeptides were normalized using PD2.5. Principal component analysis was performed in PD2.5 and Perseus v1.5.4.1 to evaluate the separation between the replicates and the sample groups. Heatmaps were prepared in Perseus v1.5.4.1 with k-means clustering and Euclidean distance. Further Cluster analysis was performed using Variance sensitive fuzzy clustering (VSClust) (http://computproteomics.bmb.sdu.dk:8192/app/VSClust) (https://doi.org/10.1093/bioinformatics/bty224). Results from the cluster analysis were further evaluated using Panther (https://pantherdb.org) against the background of all detected peptides with FDR<0.01 cut-off to identify enriched GO annotations.

### Immunoblotting

Cell on culture plates were washed twice in ice-cold PBS and lysed with modified RIPA buffer (mRIPA: 1% Triton-X, 0.1% SDS, 150 mM NaCl, 50 mM Tris pH 7.4, 2 mM EDTA, 12 mM sodium deoxycholate) freshly supplemented immediately with protease (Thermo Fisher, 78430) and phosphatase (Sigma-Aldrich, 4906845001) inhibitors. The protein concentration of the samples was measured using the Quick Start Bradford Dye Reagent (Bio-Rad, 5000205). Between 5-20 μg of total protein per sample were resolved by SDS-PAGE (4-20% gels) and transferred to Amersham Hybond P 0.45 PVDF blotting membranes (GE Healthcare, 10600023). Membranes were blocked overnight at 4°C in 5% skim milk powder or 5% BSA in PBS 0.1% Tween when working with phospho-specific antibodies, and then incubated with primary antibodies overnight at 4 °C in 5% skim milk powder or 5% BSA in PBS 0.1% Tween. HRP-conjugated goat anti-rabbit (Dako, P0448, 1:3000) and rabbit anti-mouse (Dako P0161, 1:3000) secondary antibodies were incubated for ∼1 hr at room temperature. The blots were developed with ECL Prime enhanced chemiluminescent detection reagent (GE Healthcare, RPN2232) and imaged using a Gel DocXR+ system (BioRad).

### Extracellular glucose and lactate measurements

Selected cell lines were grown on 24-well plates and fed with 0.6 mL (high) or 0.3 mL (low) of fresh medium. After 24 hours media were collected and sent for glucose consumption/lactate production analysis at the Core Biochemical Assay Laboratory (Addenbrooke’s Hospital, Cambridge). Fresh unused medium was used as a baseline. Medium glucose was measured by modified hexokinase-glucose-6-phosphate dehydrogenase method (assay DF30, Siemens Healthcare Diagnostics). Medium lactate was measured by modified Marbach and Weil Method (assay DF16, Siemens Healthcare Diagnostics).

---

**Figure S1.**
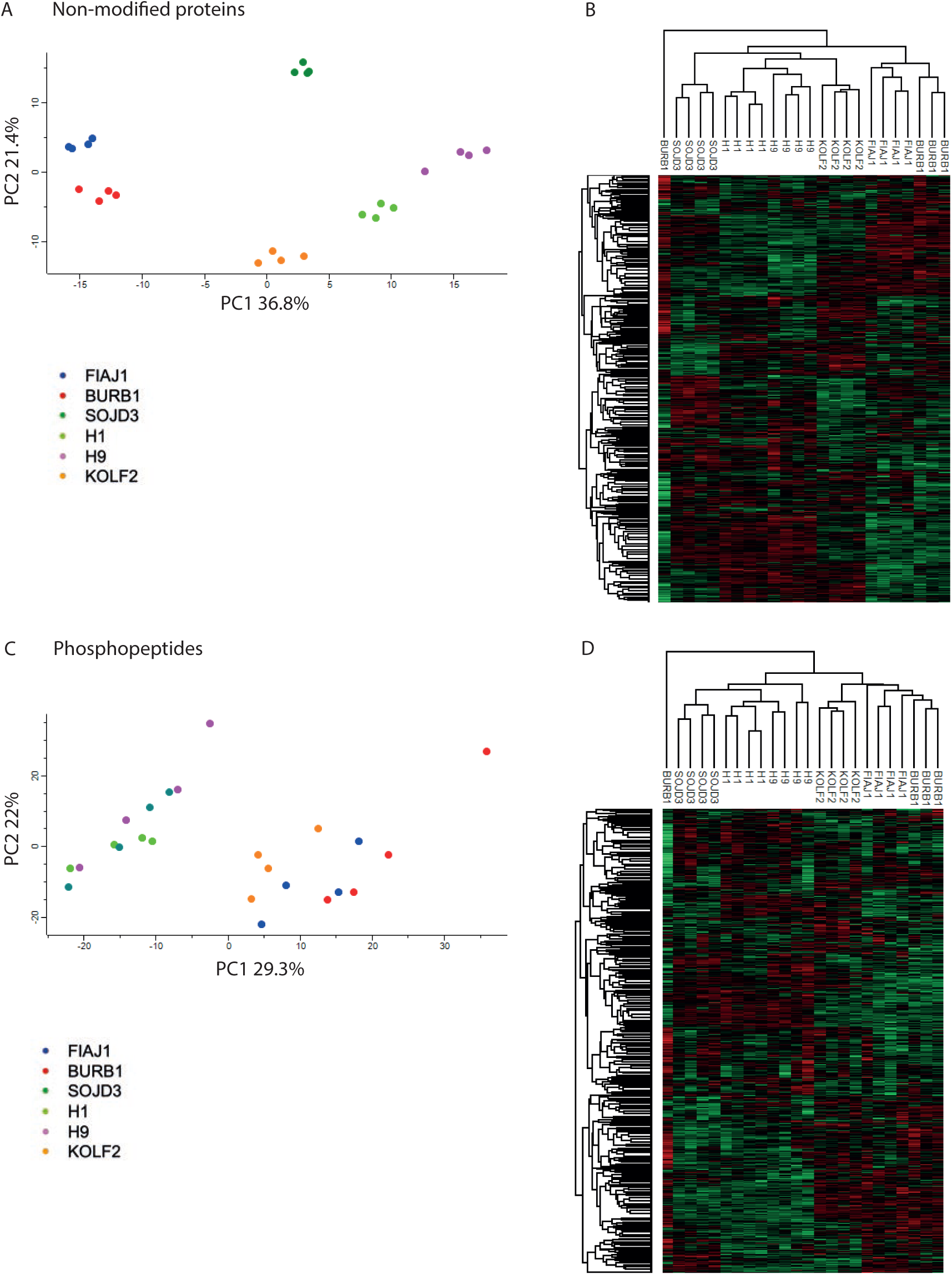
A – principal component analysis based on 5,798 non-modified proteins detected, plot shows PC1 vs PC2, B - heat map of the relative expression levels of the non-modified proteins, C - principal component analysis based on 12,143 phosphopeptides detected, plot shows PC1 vs PC2, D - heat map of the relative expression levels of phosphopeptides.

**Figure S2.**
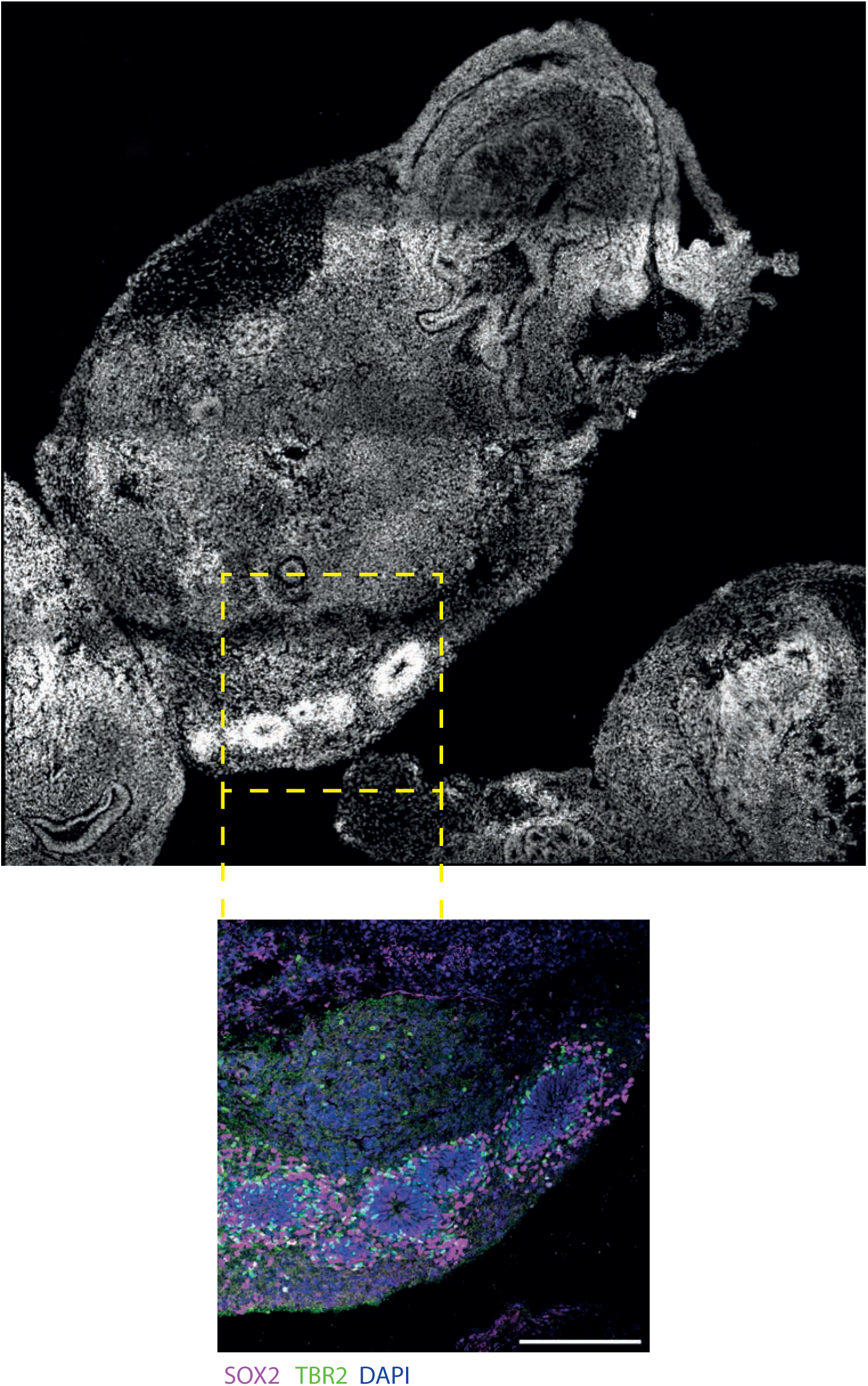
Whole sample overview of the image presented in Figure 2I, main image stained with DAPI, inset: blue – DAPI, magenta – SOX2, green – TBR2.

**Figure S3.**
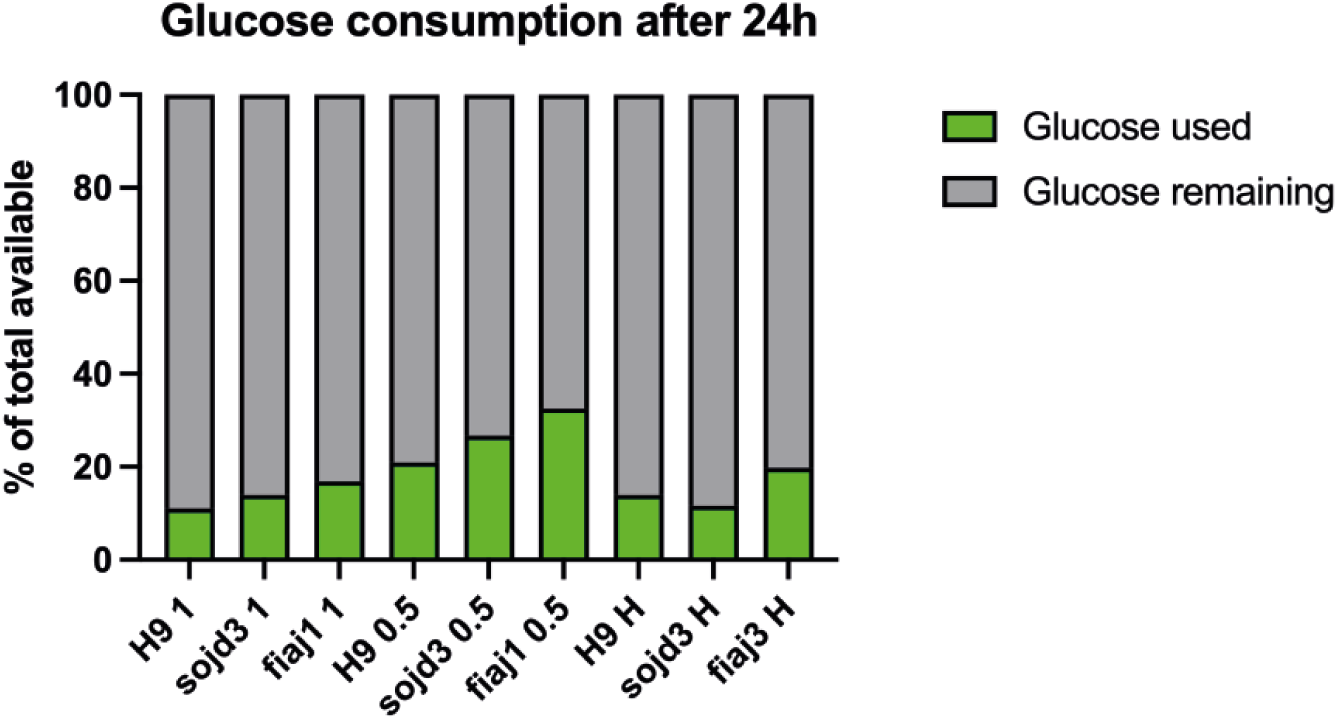
Glucose consumption after 24h, refers to Figure 3A.

## References

1. Thomson, J. A. et al. Embryonic Stem Cell Lines Derived from Human Blastocysts. Sci. New Ser. 282, 1145–1147 (1998).

2. Wataya, T. et al. Minimization of exogenous signals in ES cell culture induces rostral hypothalamic differentiation. Proc. Natl. Acad. Sci. 105, 11796–11801 (2008).

3. Ying, Q.-L., Stavridis, M., Griffiths, D., Li, M. & Smith, A. Conversion of embryonic stem cells into neuroectodermal precursors in adherent monoculture. Nat. Biotechnol. 21, 183–186 (2003).

4. Lippmann, E. S., Estevez-Silva, M. C. & Ashton, R. S. Defined Human Pluripotent Stem Cell Culture Enables Highly Efficient Neuroepithelium Derivation Without Small Molecule Inhibitors. Stem Cells 32, 1032–1042 (2014).

5. Takahashi, K. & Yamanaka, S. Induction of Pluripotent Stem Cells from Mouse Embryonic and Adult Fibroblast Cultures by Defined Factors. Cell 126, 663–676 (2006).

6. Kilpinen, H. et al. Common genetic variation drives molecular heterogeneity in human iPSCs. Nature 546, 370–375 (2017).

7. Lancaster, M. A. et al. Cerebral organoids model human brain development and microcephaly. Nature 501, 373–379 (2013).

8. Renner, M. et al. Self-organized developmental patterning and differentiation in cerebral organoids. EMBO J. 36, 1316–1329 (2017).

9. Velasco, S. et al. Individual brain organoids reproducibly form cell diversity of the human cerebral cortex. Nature 570, 523–527 (2019).

10. Camp, J. G. et al. Human cerebral organoids recapitulate gene expression programs of fetal neocortex development. Proc. Natl. Acad. Sci. 112, 15672–15677 (2015).

11. Pollen, A. A. et al. Establishing Cerebral Organoids as Models of Human-Specific Brain Evolution. Cell 176, 743–756.e17 (2019).

12. Benito-Kwiecinski, S. et al. An early cell shape transition drives evolutionary expansion of the human forebrain. Cell 184, 2084–2102.e19 (2021).

13. Lancaster, M. A. & Knoblich, J. A. Generation of cerebral organoids from human pluripotent stem cells. Nat. Protoc. 9, 2329–2340 (2014).

14. Watanabe, M. et al. TGFβ superfamily signaling regulates the state of human stem cell pluripotency and capacity to create well-structured telencephalic organoids. Stem Cell Rep. 17, 2220–2238 (2022).

15. Ideno, H. et al. Human PSCs determine the competency of cerebral organoid differentiation via FGF signaling and epigenetic mechanisms. iScience 25, 105140 (2022).

16. Glass, M. R. et al. Cross-site reproducibility of human cortical organoids reveals consistent cell type composition and architecture. Stem Cell Rep. 19, 1351–1367 (2024).

17. Sandoval, S. O. et al. Rigor and reproducibility in human brain organoid research: Where we are and where we need to go. Stem Cell Rep. 19, 796–816 (2024).

18. Merkle, F. T. et al. Whole-genome analysis of human embryonic stem cells enables rational line selection based on genetic variation. Cell Stem Cell 29, 472–486.e7 (2022).

19. Pantazis, C. B. et al. A reference human induced pluripotent stem cell line for large-scale collaborative studies. Cell Stem Cell 29, 1685–1702.e22 (2022).

20. Puigdevall, P., Jerber, J., Danecek, P., Castellano, S. & Kilpinen, H. Somatic mutations alter the differentiation outcomes of iPSC-derived neurons. Cell Genomics 3, 100280 (2023).

21. Andrews, P. W. et al. The consequences of recurrent genetic and epigenetic variants in human pluripotent stem cells. Cell Stem Cell 29, 1624–1636 (2022).

22. Jerber, J. et al. Population-scale single-cell RNA-seq profiling across dopaminergic neuron differentiation. Nat. Genet. 53, 304–312 (2021).

23. Chiaradia, I. et al. Tissue morphology influences the temporal program of human brain organoid development. Cell Stem Cell 30, 1351–1367.e10 (2023).

24. Närvä, E. et al. A Strong Contractile Actin Fence and Large Adhesions Direct Human Pluripotent Colony Morphology and Adhesion. Stem Cell Rep. 9, 67–76 (2017).

25. Stubb, A. et al. Superresolution architecture of cornerstone focal adhesions in human pluripotent stem cells. Nat. Commun. 10, 4756 (2019).

26. Albalushi, H. et al. Laminin 521 Stabilizes the Pluripotency Expression Pattern of Human Embryonic Stem Cells Initially Derived on Feeder Cells. Stem Cells Int. 2018, 7127042 (2018).

27. Laperle, A. et al. α-5 Laminin Synthesized by Human Pluripotent Stem Cells Promotes Self-Renewal. Stem Cell Rep. 5, 195–206 (2015).

28. Villa-Diaz, L. G., Kim, J. K., Laperle, A., Palecek, S. P. & Krebsbach, P. H. Inhibition of Focal Adhesion Kinase Signaling by Integrin α6β1 Supports Human Pluripotent Stem Cell Self-Renewal. Stem Cells 34, 1753–1764 (2016).

29. Li, X. et al. A fully defined static suspension culture system for large-scale human embryonic stem cell production. Cell Death Dis. 9, 892 (2018).

30. Miyazaki, T. et al. Recombinant human laminin isoforms can support the undifferentiated growth of human embryonic stem cells. Biochem. Biophys. Res. Commun. 375, 27–32 (2008).

31. Rodin, S. et al. Long-term self-renewal of human pluripotent stem cells on human recombinant laminin-511. Nat. Biotechnol. 28, 611–615 (2010).

32. Rosowski, K. A., Mertz, A. F., Norcross, S., Dufresne, E. R. & Horsley, V. Edges of human embryonic stem cell colonies display distinct mechanical properties and differentiation potential. Sci. Rep. 5, 14218 (2015).

33. Xue, X. et al. Mechanics-guided embryonic patterning of neuroectoderm tissue from human pluripotent stem cells. Nat. Mater. 17, 633–641 (2018).

34. Warmflash, A., Sorre, B., Etoc, F., Siggia, E. D. & Brivanlou, A. H. A method to recapitulate early embryonic spatial patterning in human embryonic stem cells. Nat. Methods 11, 847–854 (2014).

35. Deglincerti, A. et al. Self-organization of human embryonic stem cells on micropatterns. Nat. Protoc. 11, 2223–2232 (2016).

36. Vallier, L., Alexander, M. & Pedersen, R. A. Activin/Nodal and FGF pathways cooperate to maintain pluripotency of human embryonic stem cells. J. Cell Sci. 118, 4495–4509 (2005).

37. Dvorak, P. et al. Expression and Potential Role of Fibroblast Growth Factor 2 and Its Receptors in Human Embryonic Stem Cells. STEM CELLS 23, 1200–1211 (2005).

38. Chen, G. et al. Chemically defined conditions for human iPSC derivation and culture. Nat. Methods 8, 424–429 (2011).

39. Lotz, S. et al. Sustained levels of FGF2 maintain undifferentiated stem cell cultures with biweekly feeding. PloS One 8, e56289 (2013).

40. Bertucci, T. et al. Improved Protocol for Reproducible Human Cortical Organoids Reveals Early Alterations in Metabolism with *MAPT* Mutations. Preprint at 10.1101/2023.07.11.548571 (2023).

41. Varum, S. et al. Energy Metabolism in Human Pluripotent Stem Cells and Their Differentiated Counterparts. PLoS ONE 6, e20914 (2011).

42. Shetty, D. K., Kalamkar, K. P. & Inamdar, M. S. OCIAD1 Controls Electron Transport Chain Complex I Activity to Regulate Energy Metabolism in Human Pluripotent Stem Cells. Stem Cell Rep. 11, 128–141 (2018).

43. Zhang, J., Nuebel, E., Daley, G. Q., Koehler, C. M. & Teitell, M. A. Metabolic Regulation in Pluripotent Stem Cells during Reprogramming and Self-Renewal. Cell Stem Cell 11, 589–595 (2012).

44. Zhang, J. et al. UCP2 regulates energy metabolism and differentiation potential of human pluripotent stem cells: UCP2 regulates hPSC metabolism and differentiation. EMBO J. 30, 4860– 4873 (2011).

45. Wang, L. et al. Fatty acid synthesis is critical for stem cell pluripotency via promoting mitochondrial fission. EMBO J. 36, 1330–1347 (2017).

46. Moussaieff, A. et al. Glycolysis-Mediated Changes in Acetyl-CoA and Histone Acetylation Control the Early Differentiation of Embryonic Stem Cells. Cell Metab. 21, 392–402 (2015).

47. Ren, Z. et al. Insulin Promotes Mitochondrial Respiration and Survival through PI3K/AKT/GSK3 Pathway in Human Embryonic Stem Cells. Stem Cell Rep. 15, 1362–1376 (2020).

48. Song, C. et al. Elevated Exogenous Pyruvate Potentiates Mesodermal Differentiation through Metabolic Modulation and AMPK/mTOR Pathway in Human Embryonic Stem Cells. Stem Cell Rep. 13, 338–351 (2019).

49. Crespo, F. L., Sobrado, V. R., Gomez, L., Cervera, A. M. & McCreath, K. J. Mitochondrial reactive oxygen species mediate cardiomyocyte formation from embryonic stem cells in high glucose. Stem Cells Dayt. Ohio 28, 1132–1142 (2010).

50. Cornacchia, D. et al. Lipid Deprivation Induces a Stable, Naive-to-Primed Intermediate State of Pluripotency in Human PSCs. Cell Stem Cell 25, 120–136.e10 (2019).

51. Tan, J. et al. Oxygen is a critical regulator of cellular metabolism and function in cell culture.

52. Luengo, A. et al. Increased demand for NAD+ relative to ATP drives aerobic glycolysis. Mol. Cell 81, 691–707.e6 (2021).

53. Lees, J. G., Gardner, D. K. & Harvey, A. J. Nicotinamide adenine dinucleotide induces a bivalent metabolism and maintains pluripotency in human embryonic stem cells. Stem Cells 38, 624–638 (2020).

54. Singh, A. M. et al. Signaling Network Crosstalk in Human Pluripotent Cells: A Smad2/3- Regulated Switch that Controls the Balance between Self-Renewal and Differentiation. Cell Stem Cell 10, 312–326 (2012).

55. Ding, V. M. Y. et al. Tyrosine phosphorylation profiling in FGF-2 stimulated human embryonic stem cells. PloS One 6, e17538 (2011).

56. Wang, L. et al. Self-renewal of human embryonic stem cells requires insulin-like growth factor-1 receptor and ERBB2 receptor signaling. Blood 110, 4111–4119 (2007).

57. Smith, A. Formative pluripotency: the executive phase in a developmental continuum. Development 144, 365–373 (2017).

58. Andrews, P. W. & Gokhale, P. J. A short history of pluripotent stem cells markers. Stem Cell Rep. 19, 1–10 (2024).

59. Pagliaro, A. et al. Temporal morphogen gradient-driven neural induction shapes single expanded neuroepithelium brain organoids with enhanced cortical identity. Nat. Commun. 14, 7361 (2023).

60. Onuma, Y. et al. A Stable Chimeric Fibroblast Growth Factor (FGF) Can Successfully Replace Basic FGF in Human Pluripotent Stem Cell Culture. PLOS ONE 10, e0118931 (2015).

61. Dvorak, P. et al. Computer-assisted engineering of hyperstable fibroblast growth factor 2. Biotechnol. Bioeng. 115, 850–862 (2018).

62. Rodin, S. et al. Clonal culturing of human embryonic stem cells on laminin-521/E-cadherin matrix in defined and xeno-free environment. Nat. Commun. 5, 3195 (2014).

63. Hughes, C. S., Postovit, L. M. & Lajoie, G. A. Matrigel: A complex protein mixture required for optimal growth of cell culture. PROTEOMICS 10, 1886–1890 (2010).

64. Meng, Y. et al. Characterization of integrin engagement during defined human embryonic stem cell culture. FASEB J. 24, 1056–1065 (2010).

65. Giandomenico, S. L., Sutcliffe, M. & Lancaster, M. A. Generation and long-term culture of advanced cerebral organoids for studying later stages of neural development. Nat. Protoc. 16, 579–602 (2021).

66. Phanstiel, D. H. et al. Proteomic and phosphoproteomic comparison of human ES and iPS cells. Nat. Methods 8, 821–827 (2011).

67. Munoz, J. et al. The quantitative proteomes of human-induced pluripotent stem cells and embryonic stem cells. Mol. Syst. Biol. 7, (2011).

68. Yamamoto, T., Arita, M., Kuroda, H., Suzuki, T. & Kawamata, S. Improving the differentiation potential of pluripotent stem cells by optimizing culture conditions. Sci. Rep. 12, 14147 (2022).

69. Ji, A.-R. et al. Reactive oxygen species enhance differentiation of human embryonic stem cells into mesendodermal lineage. Exp. Mol. Med. 42, 175 (2010).

70. Mirauta, B. A. et al. Population-scale proteome variation in human induced pluripotent stem cells. eLife 9, (2020).

71. Brenes, A. J., et al. Proteomic and functional comparison between human induced and embryonic stem cells. Preprint at 10.7554/elife.92025.1 (2024).

72. Milet, C. & Monsoro-Burq, A. H. Neural crest induction at the neural plate border in vertebrates. Dev. Biol. 366, 22–33 (2012).

73. Huggins, I. J. et al. The WNT target SP5 negatively regulates WNT transcriptional programs in human pluripotent stem cells. Nat. Commun. 8, 1034 (2017).

74. Strano, A., Tuck, E., Stubbs, V. E. & Livesey, F. J. Variable Outcomes in Neural Differentiation of Human PSCs Arise from Intrinsic Differences in Developmental Signaling Pathways. Cell Rep. 31, 107732 (2020).

75. Vitillo, L., Baxter, M., Iskender, B., Whiting, P. & Kimber, S. J. Integrin-Associated Focal Adhesion Kinase Protects Human Embryonic Stem Cells from Apoptosis, Detachment, and Differentiation. Stem Cell Rep. 7, 167–176 (2016).

76. Godoy-Parejo, C., Deng, C., Liu, W. & Chen, G. Insulin Stimulates PI3K/AKT and Cell Adhesion to Promote the Survival of Individualized Human Embryonic Stem Cells. Stem Cells Dayt. Ohio 37, 1030–1041 (2019).

77. Chetty, S. et al. A Src inhibitor regulates the cell cycle of human pluripotent stem cells and improves directed differentiation. J. Cell Biol. 210, 1257–1268 (2015).

78. Lian, X. et al. A small molecule inhibitor of SRC family kinases promotes simple epithelial differentiation of human pluripotent stem cells. PloS One 8, e60016 (2013).

79. Vallier, L., Alexander, M. & Pedersen, R. A. Activin/Nodal and FGF pathways cooperate to maintain pluripotency of human embryonic stem cells. J. Cell Sci. 118, 4495–4509 (2005).

80. Martyn, I., Brivanlou, A. H. & Siggia, E. D. A wave of WNT signaling balanced by secreted inhibitors controls primitive streak formation in micropattern colonies of human embryonic stem cells. Dev. Camb. Engl. 146, dev172791 (2019).

81. Chambers, S. M. et al. Highly efficient neural conversion of human ES and iPS cells by dual inhibition of SMAD signaling. Nat. Biotechnol. 27, 275–280 (2009).

82. Yu, Y. et al. Inhibition of EZH2 Promotes Human Embryonic Stem Cell Differentiation into Mesoderm by Reducing H3K27me3. Stem Cell Rep. 9, 752–761 (2017).

83. Greber, B. et al. FGF signalling inhibits neural induction in human embryonic stem cells. EMBO J. 30, 4874–4884 (2011).

84. TeSlaa, T. et al. α-Ketoglutarate Accelerates the Initial Differentiation of Primed Human Pluripotent Stem Cells. Cell Metab. 24, 485–493 (2016).

85. Madsen, R. R. et al. NODAL/TGFβ signalling mediates the self-sustained stemness induced by PIK3CAH1047R homozygosity in pluripotent stem cells. Dis. Model. Mech. 14, dmm048298 (2021).

86. Larsen, M. R., Thingholm, T. E., Jensen, O. N., Roepstorff, P. & Jørgensen, T. J. D. Highly Selective Enrichment of Phosphorylated Peptides from Peptide Mixtures Using Titanium Dioxide Microcolumns. Mol. Cell. Proteomics 4, 873–886 (2005).

87. Schweppe, D. K. et al. Full-Featured, Real-Time Database Searching Platform Enables Fast and Accurate Multiplexed Quantitative Proteomics. J. Proteome Res. 19, 2026–2034 (2020).

